# A novel enrichment strategy reveals unprecedented number of novel transcription start sites at single base resolution in a model prokaryote and the gut microbiome

**DOI:** 10.1101/034785

**Authors:** Laurence Ettwiller, John Buswell, Erbay Yigit, Ira Schildkraut

## Abstract

We have developed Cappable-seq that specifically captures primary RNA transcripts by enzymatically modifying the 5’ triphosphorylated end of RNA with a selectable tag. We first applied Cappable-seq to *E. coli*, achieving up to 50 fold enrichment of primary transcripts and identifying an unprecedented 16539 transcription start sites (TSS) genome-wide at single base resolution. We also applied Cappable-seq to a mouse cecum sample and for the first time identified TSS in a microbiome. Furthermore, Cappable-seq universally depletes ribosomal RNA and reduces the complexity of the transcriptome to a single quantifiable tag per TSS enabling digital profiling of gene expression in any microbiome.

## Background

High-throughput cDNA sequencing has emerged as a powerful tool to globally assess the transcriptional state of cells. However, post-transcriptional processing and modification events add layers of complexity to transcriptomes that are typically not revealed by standard RNA-seq technologies. For example, processed ribosomal RNA (rRNA) typically constitutes 95% of the total RNA in prokaryotes with only a minority of the RNA corresponding to protein coding transcripts [1]. Such RNA processing confounds the identification of key transcriptional events such as the start and end of transcription and, more generally, the original composition of primary transcripts. Thus, being able to decouple the primary transcriptome from processed RNA is key to determining the association between the regulatory state of the genome and its phenotypic outcome. Identifying the primary transcriptome depends on the ability to distinguish the initiating 5’ nucleotide incorporated by the RNA polymerase from all the other 5 ‘ ends that arise due to processing. This distinction is not feasible with the currently available methods.

Here we present a significant advance in transcriptomics to directly and universally target the first nucleotide that has been incorporated by the RNA polymerase upon initiation of transcription. This nucleotide marks the transcription start site on the genomic sequence. Our strategy consists of enzymatically labeling, with a biotin derivative, transcripts that have retained their original initiating 5’ nucleotide. Only transcripts that have an intact 5’ triphosphorylated (or 5’ diphosphate) end are biotinylated and isolated from the in-vivo processed RNA. We refer to enzymatic labeling of the 5’ triphosphorylated end of RNA and subsequent enrichment and high-throughput sequencing as Cappable-seq.

Cappable-seq has a broad range of applications, offering the ability to investigate the triphosphorylated population of RNA molecules that would otherwise be masked by the overwhelming majority of their processed counterparts. By accurately anchoring the origin of the transcript to single base specific position on the genome, Cappable-seq reduces sequence complexity to a unique tag per transcript. The identification of the TSS to single base resolution enables the association between the regulatory state of a genome and its transcriptome. Thus, changes in transcription factor binding profiles and/or epigenetic states, notably at promoters, can be associated with changes in transcription by quantifying transcription start site (TSS) usage. While various methods for determining prokaryotic TSS have been developed, all of them attempt to circumvent the inability to directly capture the 5’ triphosphorylated ends. The most widely used method, TEX relies on eliminating the processed transcripts by treating RNA samples with Xrn1 exonuclease. This exonuclease preferentially degrades RNAs containing a 5’ monophosphate, therefore resulting in an apparent enrichment of primary transcripts containing 5’-triphosphates [1–7]. To increase the specificity of the TEX method, a control non-Xrn1 treated library is subtracted from the TEX library. This method is referred to as differential RNA-seq (dRNA-seq). Nevertheless, TEX and dRNA-seq have been previously reported to suffer from a lack of specificity for TSS [8–10].

As a proof of concept, we applied Cappable-seq for the precise determination of TSS genome-wide in *E. coli*. Cappable-seq was performed on total RNA and a remarkable number of 16359 TSS at single base resolution were found. We show that Cappable-seq is highly specific to triphosphorylated RNA characteristic of TSS. Compared to RNA-seq, Cappable-seq reduces the complexity of the transcriptome, enabling digital profiling of gene expression. Processed ribosomal RNA are also reduced from an overwhelming majority of total RNA to only 3%, allowing a deeper sequencing of the informative transcriptome at lower cost. By applying Cappable-seq to a mouse cecum sample, we demonstrate for the first time, identification of TSS from a microbiome. Cappable-seq applied to mouse cecum microbiome identified TSS in species from different bacterial phyla. We found novel promoter consensus regions in all phyla analyzed and leaderless transcripts that account for 10 to 15% of identified TSS in some species of the microbiome such as *Akkermansia muciniphila* and *Bifidobacterium pseudolongum*. Ribosomal RNA represents less than 5% of RNA for the majority of species analyzed suggesting that most of the sequences represent TSS of protein coding transcripts. Thus, this methodology provides a unique solution for TSS determination and digital profiling of gene expression of microbiomes while universally removing the contaminating ribosomal RNA that constitute the major cost burden of transcriptomes and meta-transcriptomes.

## Results

### Cappable-seq captures the triphosphorylated RNA and enriches for primary transcripts

Cappable-seq isolates the primary transcripts by enzymatically capping of the 5 ‘ triphosphorylated RNA with a biotinylated GTP using vaccinia capping enzyme (VCE). For this purpose, we screened a number of biotinylated derivatives of GTP and found that 3’ OH modifications of ribose of GTP are acceptable substrates for VCE. The biochemistry of capping and decapping are presented in Supplemental Note A and Supplemental Fig. 1, 2 and 3 (All supplemental notes and figures are in Additional file 1). The reaction results in the specific labeling of 5’-di or triphosphorylated RNA ends while the 5’-monophosphorylated RNA ends characteristic of processed transcripts are not labeled (Supplemental Fig. 2 and 4). The biotinylated RNA can then be captured on streptavidin beads and isolated (Supplemental Fig. 3).

### Application of Cappable-seq to *E. coli* reveals an unprecedented number of TSS

We first applied Cappable-seq for the genome-wide identification of TSS in the model organism *E. coli* MG1655. For this, total *E. coli* RNA was capped with 3’-desthiobiotin-TEG-guanosine 5’ triphosphate (DTBGTP) for reversible binding to streptavidin, fragmented to an approximate size of 200 bases, captured on streptavidin beads and eluted to obtain the 5’ fragment of the primary transcripts (see method section and Figure 1A). To achieve single base resolution, a Cappable-seq library was generated by ligating 5’ and 3’ adaptors to the RNA. In this case, the labeled cap must first be removed from the RNA to allow the ligation to the 5’end. We found that RppH efficiently removes the desthiobiotinylated cap structure to leave a ligatable 5’-monophosphate RNA (Supplemental Fig. 5 and 6).

**Figure 1:**
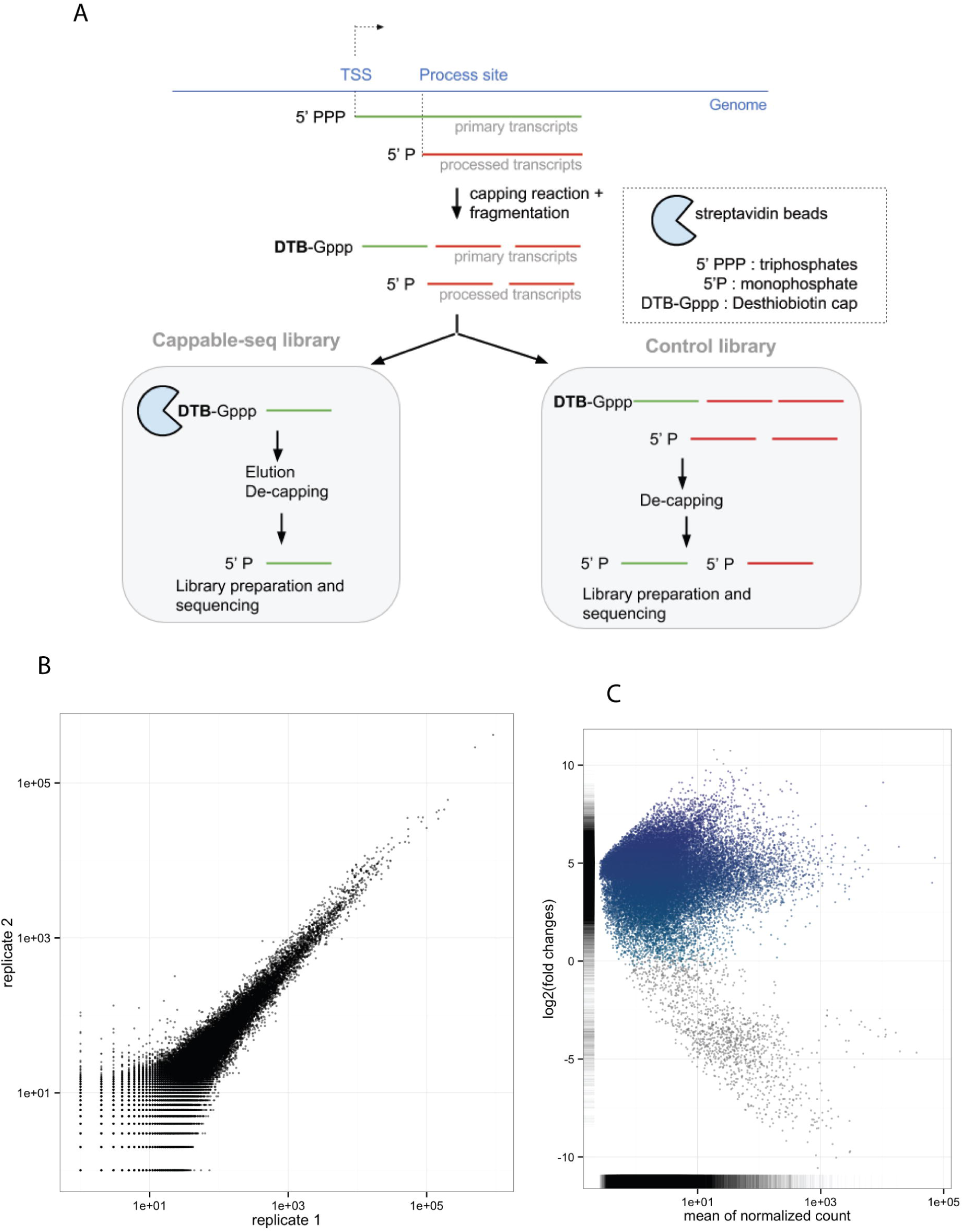
Cappable-seq pipeline for TSS identification. **A**. Schema of Cappable-seq protocol and the associated control library. **B**. Replicate analysis. The correlation coefficient between replicate 1 and replicate 2 RRS is 0.983. **C**. Enrichment score as a function of the mean of relative read score for the 36078 putative TSSs found in *E. coli* grown on minimal media. In blue are TSS that are enriched in Cappable-seq library. Grey are positions that are depleted in Cappable-seq. The removal of depleted positions eliminates 1354 spurious TSS primarily located in ribosomal loci.

A non-enriched control library was prepared using identical conditions as Cappable-seq except that the streptavidin capture step was omitted. Both libraries were sequenced using Illumina MiSeq yielding approximately 20 million single end reads. Reads were mapped to the *E. coli* genome using Bowtie2 [11]. The orientation and mapped location of the first base of the sequencing read determines the genomic position of the 5’ end of the transcript at single base resolution. The number of reads at a specific position defines the expression level of the 5’ end of the transcript. We normalized this number with the total number of mapped reads to obtain a relative read score (RRS) reflecting the strength of each TSS, thus defining a single quantifiable tag per transcript that can be used for digital gene expression profiling. A technical replicate generated using the same total *E. coli* RNA preparation resulted in a correlation coefficient of 0.983 demonstrating the high reproducibility of Cappable-seq (Figure 1B).

The ratio between the RRS from Cappable-seq and the non-enriched control libraries defines the enrichment scores with enriched positions corresponding to 5’-triphosphorylated ends characteristic of TSS and depleted positions corresponding to processed/degraded 5’ ends (see Supplemental note B in Additional file 1 and Figure 1C). To define TSS, we selected positions on the genome with a RRS of 1.5 and higher (equivalent to 20 reads or more) and found 36,078 positions satisfying this criterion. Next, we subtracted the 1354 positions that are depleted in the Cappable-seq library when compared to the non-enriched control library (method and Figure 1C). This resulted in 34724 unique positions that we define as TSS. This step reduces the number of positions by only 3.7%. As most of the false positive positions are located in ribosomal genes, the exclusion of positions located within those genes drops the false positive rate to only 1.4%. Therefore the need to sequence a non-enriched RNA library in order to calculate an enrichment score is not critical with Cappable-seq whereas a non-enriched library is required to perform dRNA-seq [7].

The accurate description of TSS in prokaryotes relies on the differentiation of the 5’-triphosphorylated end that characterizes primary transcripts from the 5’-monophosphorylated end that characterizes processed sites. Comparing the results of Cappable-seq with the results of Kim[3] and Thomason[7] demonstrates the higher specificity of Cappable-seq for 5’ triphosphate RNA (see supplemental note B and Supplemental Fig. 7). Indeed while Cappable-seq correctly calls 110 out of 111 processed sites, dRNA-seq [7] mis-annotated 40 of the processed sites as TSS (Supplemental Fig. 7B).

The higher specificity of Cappable-seq for the 5’ end of primary transcripts also has the desirable property of reducing reads mapping to rRNA from 85% of total reads to only 3% (Supplemental Fig. 7A). While some remaining reads may be background noise, we identify 26 enriched positions within rRNA genes suggesting bona-fide TSS falling within the rRNA genes (Supplemental Fig. 8).

### Genome-wide position of TSS suggests both precise and imprecise initiation of transcription

We and others have observed that many promoters initiate a low level of transcription from multiple positions closely surrounding the major initiation site for a given TSS [12]. We hypothesize that those sites may have been generated from a single promoter and thus are considered dependent. We clustered all TSS generated from a unique promoter event to one single position with the highest RRS resulting in 16359 unique positions that we define as clustered TSS (Supplemental note C and Supplemental Fig. 9A and Supplemental Table 1 in Additional file 2).

While the RNA polymerase initiates transcription at imprecise positions for about 60% of the promoters, 40% have precise positions. Interestingly, the degree of precision in the initiation site is dependent on the sequence context at TSS where the −1 and +1 positions of the TSS correspond to pyrimidine (Y) and purine (R) respectively. The −1+1 YR motif correlates with precise initiation events (Supplemental note C and Supplemental Fig. 9B).

### 41% of Cappable-seq TSS in *E. coli* are novel

To estimate how many of the TSS found by Cappable-seq are novel, we compiled a composite dataset consisting of the annotated RegulonDB TSS plus TSS derived from high throughput methodologies that have been done on *E. coli* grown in similar conditions [3, 7] and compared the 16855 TSS present in the composite dataset to the 16359 TSS determined by Cappable-seq. We identified 9600 common TSS present in both the composite dataset and Cappable-seq and 6759 TSS unique to Cappable-seq (41.3 %)(Figure 1C). The number of novel TSS that Cappable-seq identifies that have not been identified in previous studies under equivalent growth conditions is remarkable. The profile of enrichment scores (Supplemental Fig. 10A) and promoter structure (Fig. 2) is similar for both the overlapping and Cappable-seq specific sets suggesting that those novel positions are bona-fide TSS.

**Figure 2:**
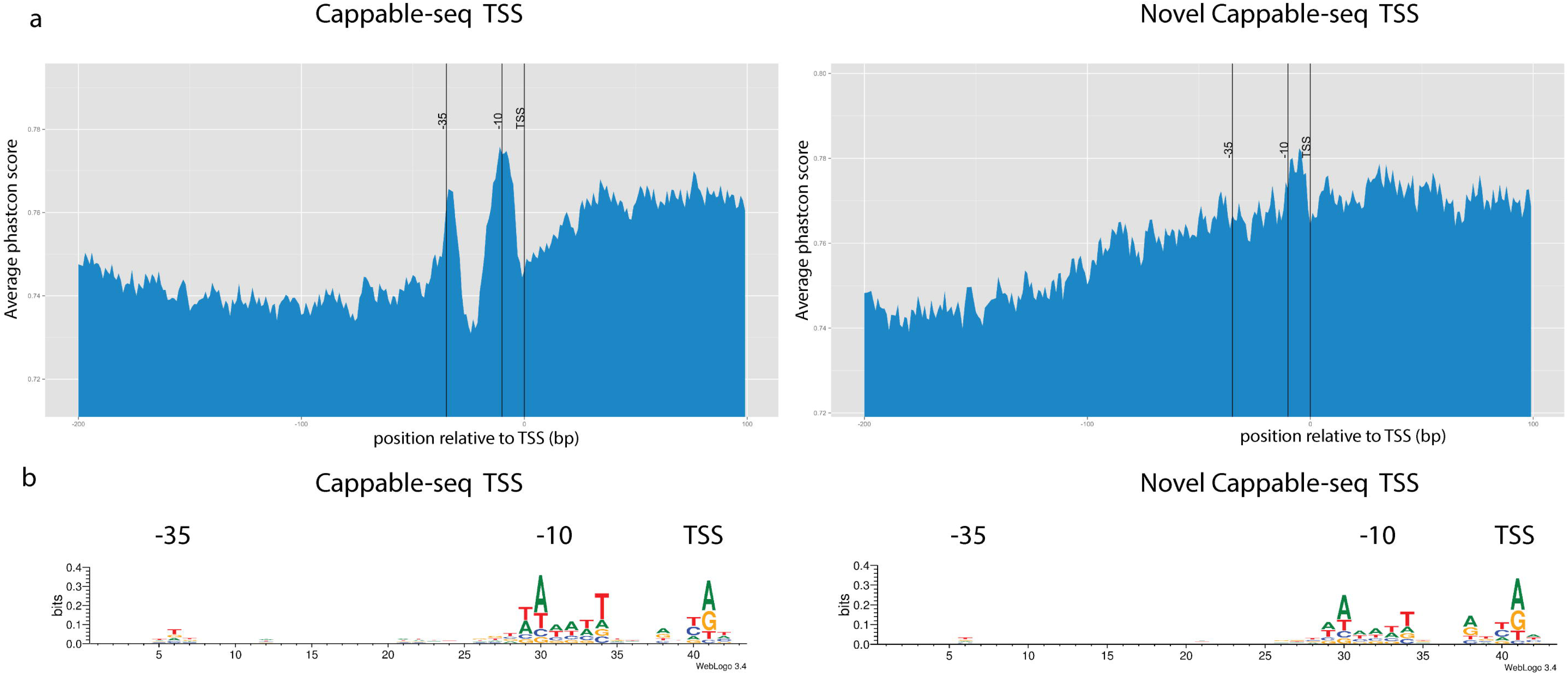
Promoter regions. Characteristics of the promoter region. **A**. The average phastcon score is plotted for each position from −100 bases upstream to +30 bases downstream of the 16359 TSS. **B**. Sequence logo upstream of all Cappable-seq TSS and novel Cappable-seq TSS.

One explanation for the high number of novel TSS with Cappable-seq is the increased sensitivity due to the higher sequencing depth, revealing novel TSS that are weakly expressed. We addressed this question by looking at the distribution of expression level for both the previously annotated and novel TSS and found a higher number of weak TSS in the Cappable-seq specific set (mean of 2.8) compared to the common set (mean of 4.9) (Supplemental Fig. 10B). Taken together, these results suggest that some novel TSS are explained by the gain of sensitivity from a high sequencing depth.

It is conceivable that an even deeper sequencing depth with Cappable-seq would reveal even more novel TSS and it is unclear at what depth this trend will cease. Such weakly expressed TSS maybe the reflection of stochastic events resulting from the transcriptional machinery occasionally initiating transcription from non-canonical promoters. This stochastic initiation would result in an increased repertoire of transcripts conferring phenotypic diversity to an otherwise genotypically identical population of cells. Analogous to the inherent mutation rate of DNA polymerases as a driver for evolution [13], we hypothesize that stochastic transcription starts may confer an evolutionary advantage.

### Upstream regions of TSS display characteristics of known *E. coli* promoters

Next, we analyzed the sequence conservation across related species and nucleotide bias upstream of the 16359 Cappable-seq TSS. To calculate the overall conservation of the flanking regions of TSS, we used the phastcon scores [14] derived from the genome-wide alignment of 10 related bacterial species including *E. coli* from UCSC (Materials and Methods). As expected, the overall conservation score increased at around 10 and 35 bp upstream of TSS and gradually increased downstream of the TSS (Figure 2A). The upstream conservation is indicative of the presence of the −10 and −35 promoter elements suggesting that a significant fraction of promoters upstream of the Cappable-seq TSS are under positive selection. The downstream conservation across the ten listed species is indicative of open reading frames likely present downstream of TSS. Nucleotide bias in the region upstream of the TSS is in accordance with sequence conservation; there is a strong bias at −10 for a motif resembling the TATAAT box and a weaker bias at −35 resembling the sigma factor 70 binding site (Figure 2B). Taken together, these results are consistent with the structure of *E. coli* promoters, particularly the sigma 70 promoters upstream of a majority of TSS.

We performed the same analysis with the 6759 novel Cappable-seq TSS and found that the regions show similar sequence bias at around −35 and −10 as that found for the entire set (Figure 2B). Interestingly, despite similar sequence bias in both novel and annotated TSS, the novel TSS show no increase of sequence conservation at −10 and −35 (Figure 2A).

We also found that Cappable-seq TSS demonstrated an 80% nucleotide preference for either A or G (Figure 3A). While this finding is in agreement with previous studies [3, 12], the preference for A or G in Cappable-seq TSS is stronger than the preference found in annotated TSS from RegulonDB [15] (60%). Interestingly, despite motif preferences at the TSS, the sequence conservation across species is not elevated suggesting there is not a strong selective pressure to conserve a specific nucleotide.

**Figure 3:**
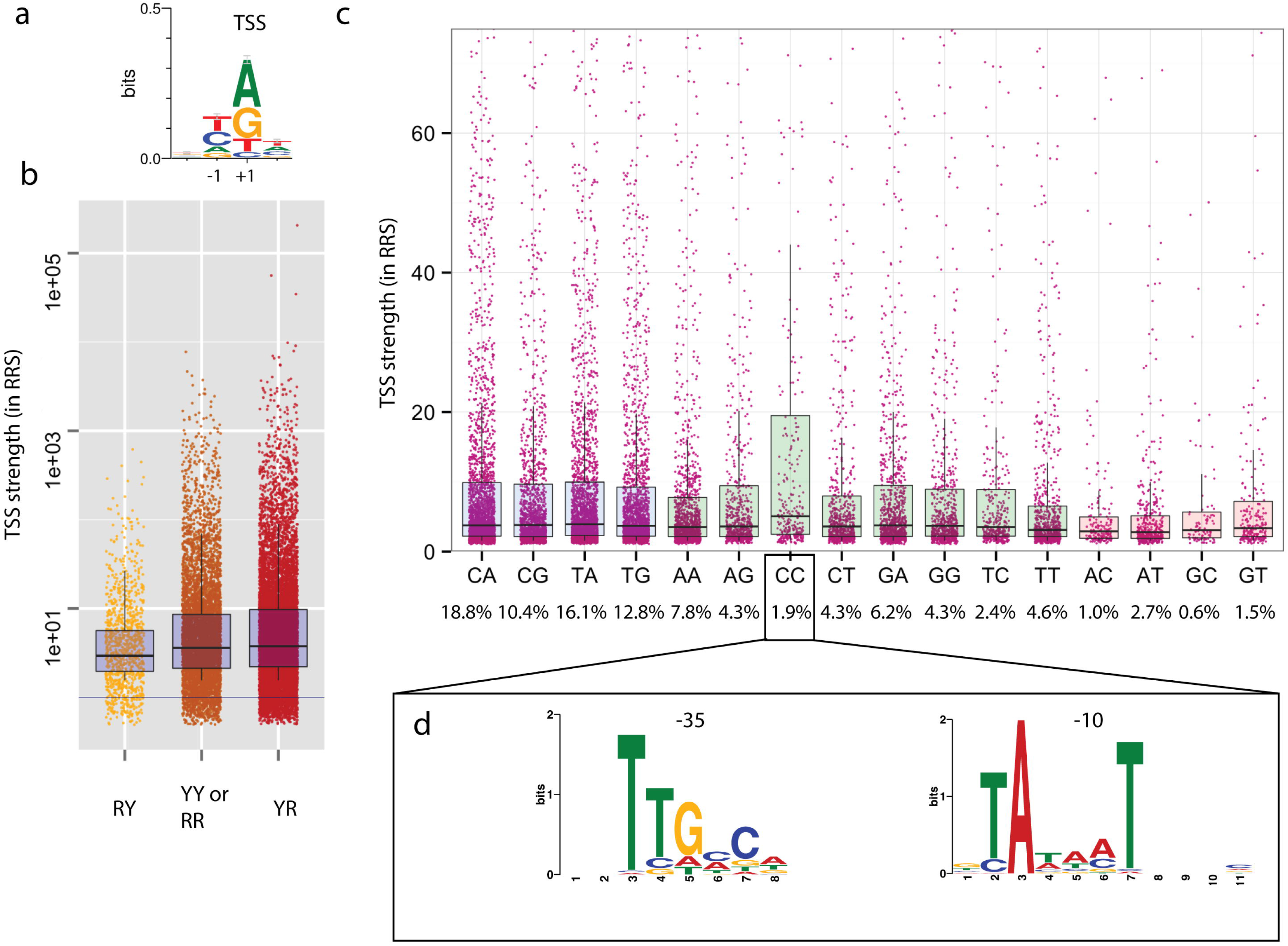
Nucleotide preference at TSS. **A**. Sequence logo of the nucleotide bias from −2 to +2 position of TSS. **B**. Distribution of the strength of the TSS (in RRS in Cappable seq) as classified according to their −1+1 configuration with R being purine (A or G) and Y being pyrimidine (C or T). **C**. Relative abundance of reads for each of the 16 possible TSS −1 +1 dinucleotides. Blue boxes are YR motifs, green boxes are YY or RR motifs and pink boxes are RY motifs. Percentages corresponds to the percentage of TSS having the aforementioned −1+1 configuration **D**. Over-represented motifs at −35 and −10 bp upstream of TSS with the −1C+1C dinucleotide configuration.

Additionally, we observed a nucleotide preference at minus 1 position with 76% of the nucleotides being pyrimidine (C or T). In summary, more than half of the TSS (57%) have a −1[CT]+1[AG] configuration with 18% of the TSS having a −1C+1A configuration and only 0.6% having the −1G+1C configuration (Figure 3C). Interestingly this pyrimidine (Y) purine (R) or “YR” configuration has been previously reported to be the preferred configuration at TSS in various prokaryotes and eukaryotes ranging from *C. elegans*, plant and human [16–18] suggesting that the YR rule is conserved across kingdoms.

There is no correlation between the −1/+1 nucleotide and the enrichment score (data not shown) suggesting that the least favored configurations (-1[AG]+1[CT]) are genuine TSS. The strength of the TSS, as defined by the RRS, has a weak correlation with the −1/+1 nucleotide configuration. Indeed, YR configuration includes the most highly expressed TSS while the RY configuration is the weakest TSS (Figure 3B). Contrasting with this notion, the −1C+1C (YY configuration) has the highest fraction of highly expressed TSS (Figure 3C) including the five most highly expressed −1C+1C TSS upstream of ribosomal genes. This observation could be the result of an alternative promoter upstream of the −1C+1C TSS. To address this question, we searched for overrepresented motifs in the 40 bases upstream of −1C+1C TSS class using MEME [19] and found the canonical TATAAT box at −10 and sigma 70 motif at −35 suggesting that the majority of the −1C+1C TSS class is a subset of TSS from the sigma 70 promoter (Figure 3D).

### Intragenic sense TSS in *E. coli* have a marked preference for the first nucleotide of codons

TSS identified by Cappable-seq that are within protein coding genes account for 63% (10741) of the total TSS with two-thirds of the intragenic TSS in the sense orientation in relation to the gene. Sense TSS tend to be located at the start of the protein coding regions while the antisense tend to be evenly distributed within the protein coding regions (Figure 4A). Intergenic TSS tend to be have higher RRS than both sense and antisense intragenic TSS, suggesting that intergenic TSS tend to be stronger (Figure 4B). There is a correlation between the strength of sense TSS and its position relative to the coding gene with stronger TSS occurring towards the 3’end of genes (Figure 4C). Leaderless transcripts account for 0.4% (82) of TSS.

**Figure 4:**
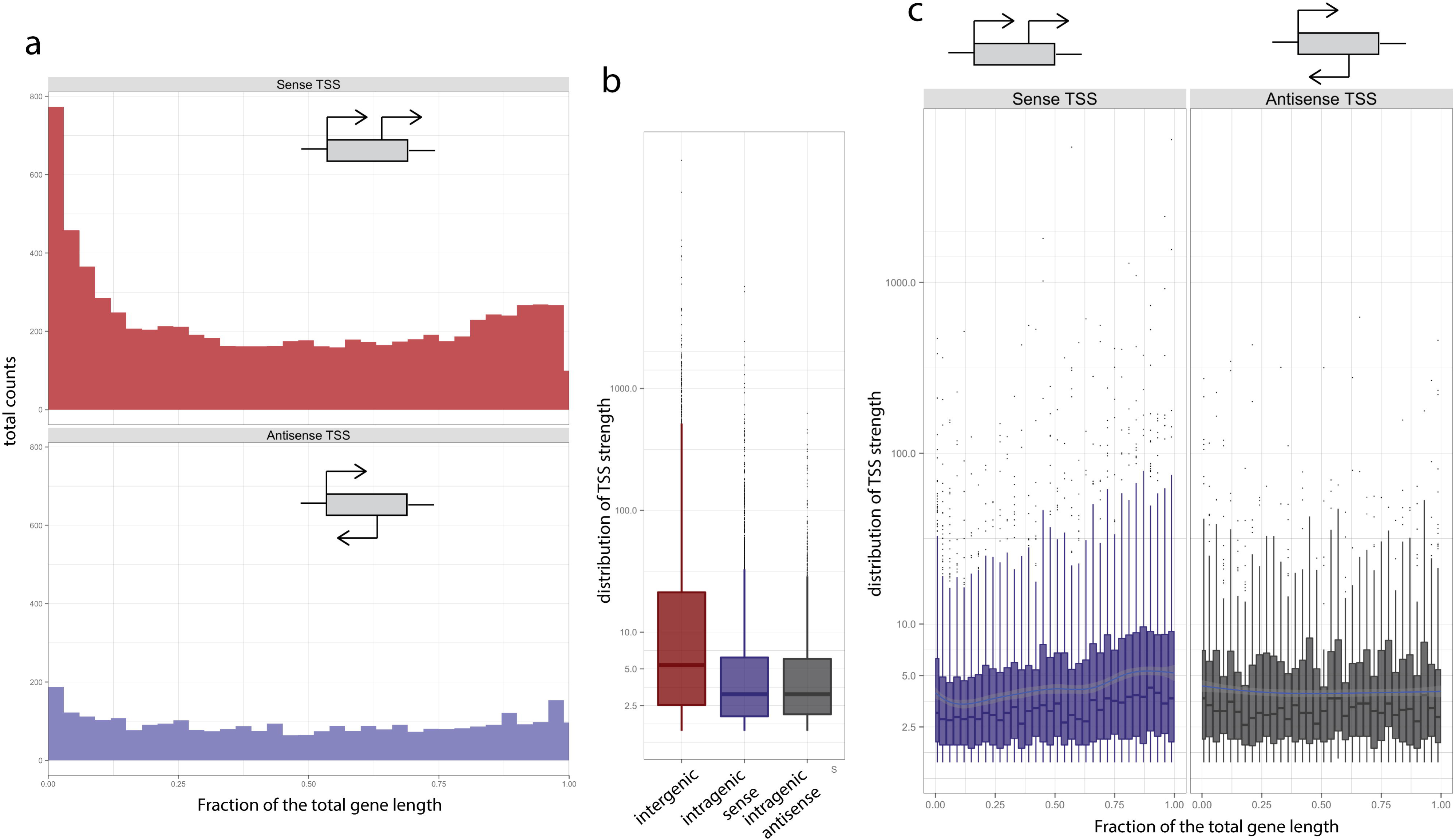
Intragenic TSS. **A**. Distribution of the number of sense and antisense intragenic TSS as a function of the position within genes. **B**. Box plot representing the distribution of the TSS strength (RRS score) for intergenic (red), sense intragenic (blue) and antisense intragenic (grey) TSS. **C**. Distribution of intragenic sense (blue) and antisense (grey) TSS strength as a function of their position within genes.

Interestingly, we found that intragenic TSS have striking positional preference relative to the nucleotide triplet that defines the reading frame. We found that 45% of the intragenic sense TSS are located in the first position of codons while only 27% of TSS are located in the second and 27% in the third position. (Figure 5A). The antisense TSS show a weaker but noticeable preference for the third position rather than the first, with 43% of TSS on the third position (Figure 5B). Sense and antisense preference is distributed throughout the protein coding gene (Figure 5A and B). This positional preference of the TSS relative to the codon may be influenced by the nucleotide frequency at codons with a higher A and G frequency at the first base of the codon. While other datasets derived from dRNA-seq experiments [7] show similar preferences, this observation has not been previously reported. Interestingly, we found 168 TSS at the first nucleotide of an internal in-frame AUG codon. Those transcripts are putative leaderless transcripts leading possibly to a truncated form of the annotated protein.

**Figure 5:**
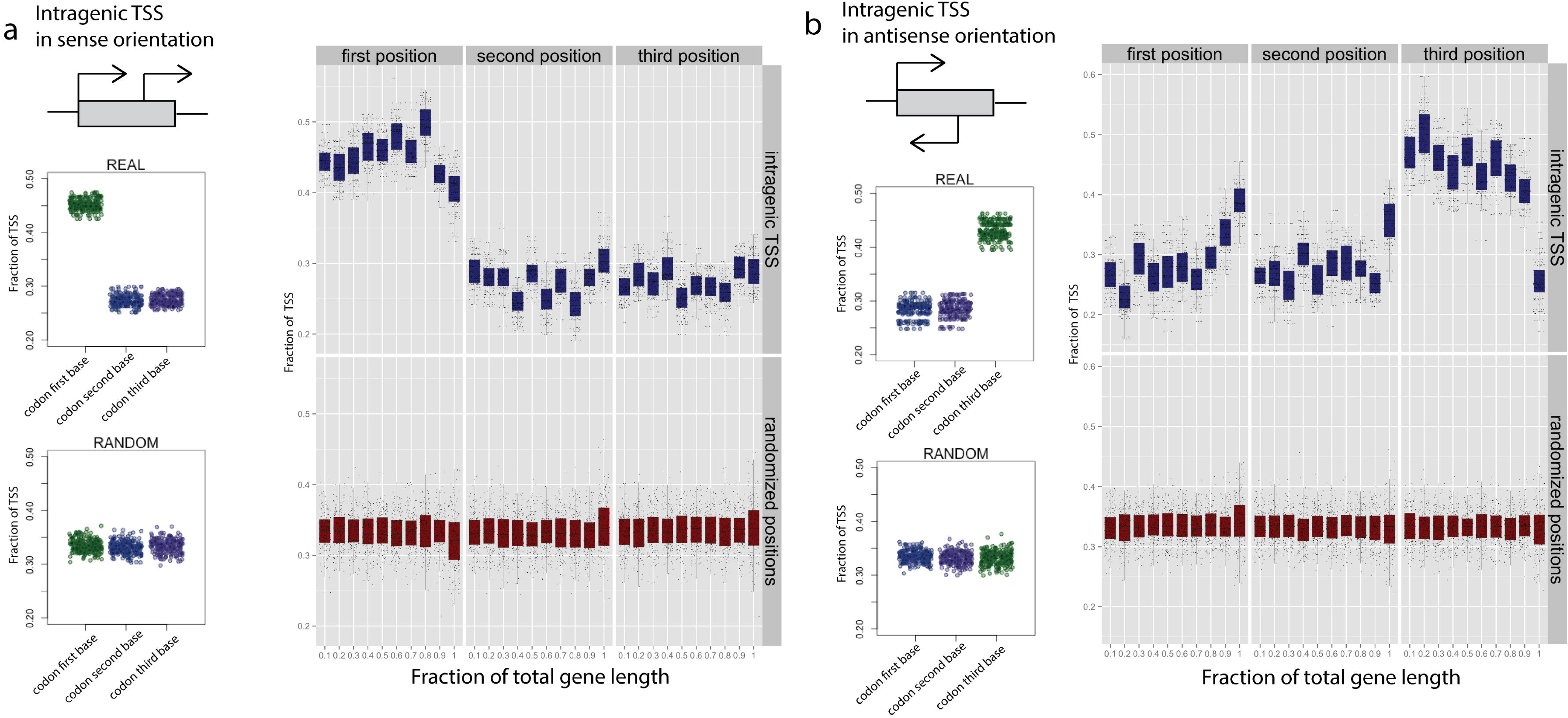
Positional preference of TSS relative to codon. Frequency of intragenic TSS relative to the first, second and third position of the codon for (**A**) the sense TSS and (**B**) the antisense TSS. Graphics on the left represent the overall frequency of TSS at each codon position across the entire gene length while the graphic on the right represent the frequency of TSS at each codon position as a function of the relative position within the coding gene (in 10 % increments of the total gene length).

### TSS from a microbiome

To demonstrate the applicability of our methodology on a complex mixture of bacteria, we applied Cappable-seq to two C57 female mice cecum microbiomes (Materials and Methods). Reads were mapped to the bacterial genomes from NCBI and species with more than 300 identified clustered TSS were considered candidates and the species with the highest number of clustered TSS in each phylum were further analyzed. For all species, we found that the majority of the reads mapped in either intergenic regions or in protein coding genes in accordance with the biology of transcription start sites (Figure 6D). Accordingly, reads mapping to rRNA and transfer RNA (tRNA) account for less than 10 % of mappable reads in *Lactobacillus johnsonii, Akkermansia muciniphila* and *Lachnospiraceae bacterium*. We hypothesize that the higher fraction of rRNA reads in *Bifidobacterium pseudolongum* (around 30%) is due to the high level of rRNA sequence conservation leading to the spurious mapping of rRNA sequence originating from other related species of *Bifidobacterium*. Taken together these data suggest that Cappable-seq depletes processed transcripts such as rRNA and tRNA from microbiomes total RNA with the same efficiency as observed in *E. coli*. Next, we derived a set of highly confident TSS per species and identified sequence bias in regions flanking those TSS. In agreement with promoter organization/structure in bacteria, we found a strong sequence bias at 35 bases and 10 bases upstream of the TSS for all analyzed species (Figure 6B) indicative of the −35 element and the TATAAT box respectively. Furthermore, the YR motif at position −1+1 can be identified in all cases, reinforcing the universality of the YR motif for TSS. Beyond the biological significance of these finding, these results shows that the specificity of Cappable-seq for TSS in a microbiome is similar to the specificity for TSS in *E. coli*. Interestingly, two of the four species analyzed (*Akkermansia muciniphila* and *Bifidobacterium pseudolongum)* show 10% and 15% of the TSS located at the start of the annotated protein coding genes which is a signature of leaderless transcripts (Figure 6C). For comparison, *E. coli* shows only 0.3% leaderless TSS. This result is in agreement with previous computational predictions [20] suggesting that leaderless transcripts are widespread in a variety of bacteria. Finally, we challenged the reproducibility of Cappable-seq in a microbiome by analyzing the TSS positions and strength (RRS) in two biological replicates from two different mice and found good reproducibility in both qualitative and quantitative (correlation coefficient = 0.81) measurements of TSS (Figure 6A[21, 22]-E). Summing up, the collective results obtained using Cappable-seq on the mouse gut microbiome demonstrates the utility and reproducibility of Cappable-seq for meta-transcriptome analysis.

**Figure 6:**
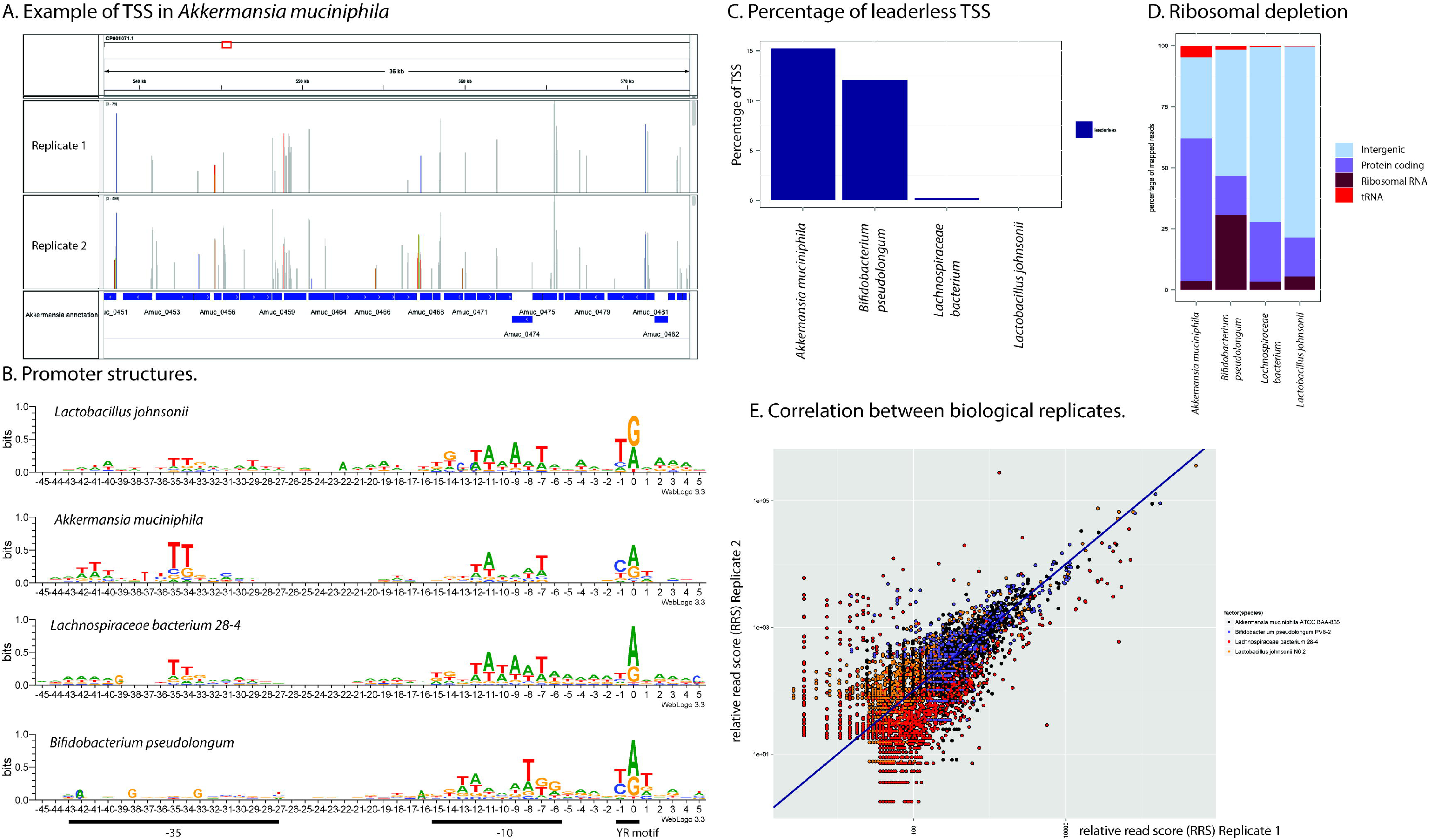
TSS of mouse gut microbiome. Analysis of TSS for four representative species across four phyla of bacteria. A. IGV display of read distribution in *Akkermansia muciniphila* in both biological replicates. B. Promoter structures in all four species generated with Weblogo (for Biological replicate 1). The X axis represent the distance away from the TSS found by Cappable-seq. Y axis represent the amount of information present at every position in the sequence, measured in bits. C. Percentage of leaderless TSS in replicate 1. D. Read genomic distribution for replicate 1. E. The correlation coefficient of relative read score (RRS) of TSS in the four representative species between the two biological replicate (two mouse gut microbiome) is 0.81.

### Discussion

Cappable-seq is a novel method that enables direct modification and identification of the triphosphorylated RNA characteristic of primary transcripts. In this study, we demonstrate the ability of Cappable-seq to determine TSS at one base resolution genome-wide in *E. coli* by pairing Cappable-seq with direct 5 ‘ ligation of sequencing adaptors to the RNA. Despite being a very different approach for determining TSS, the results are consistent with the established methodologies. Indeed, a large fraction (59 %) of the TSS found in *E. coli* by Cappable-seq is coincident with annotated TSS. Conversely, 44 % of the annotated TSS from the composite dataset are not identified by Cappable-seq. We believe that the unidentified TSS may be due to different growth conditions as well as false positives in the composite dataset. Indeed the composite dataset includes data derived from genome-wide dRNA-seq / TEX technologies and RegulonDB which is a curated dataset of TSS derived by an array of different technologies one of them being individual primer extension reactions. We and others have shown that dRNA-seq and TEX lack specificity [8, 9] for TSS and primer extension will identify any 5’ end of RNA regardless of origin. Thus, some of the genomic positions identified as TSS in the composite dataset may be false positives. In contrast, we show that Cappable-seq discriminates the 5’ triphosphate end characteristic of initiating 5 ‘ triphosphorylated nucleotide incorporated by the RNA polymerases from the processed 5’ monophosphate RNAs. This property can also be used to determine processed sites, rather than TSS, by identifying the depleted positions in Cappable-seq. We applied this analysis to our data and found approximately 3000 processed sites in the *E. coli* genome (data not shown). This assessment of processed sites is analogous to the method used by Romero [9] where the libraries have been prepared with and without tobacco acid pyrophosphatase.

Cappable-seq performs well when applied to a mouse gut microbiome and provides for the first time a solution for TSS determination in complex microbiome population. Thus, Cappable-seq can be used to derive sets of quantitative markers from which association to diseases or direct perturbation of the microbiome can be made. This technology can greatly facilitate metagenome-wide association studies by providing a signature profile of the microbiome functional state. In prokaryotes, the DTBGTP enrichment on unfragmented RNA can be used for whole transcriptome analysis by RNA-seq effectively removing rRNA. Such rRNA depletion is ideally suited for microbiome studies as it should universally remove rRNA and most contaminating eukaryotic host RNA leaving prokaryotic transcripts intact.

Applying Cappable-seq directly to total eukaryotic RNA would reveal the triphosphorylated transcriptome derived from Pol I and III RNA polymerases and identify the TSS of these transcripts. Eukaryotic pol II transcripts differ from Pol I and III transcripts by virtue of their 5’ G cap. Thus, the removal of the G cap with a decapping enzyme, which leaves a recappable 5’ diphosphate at the 5’end of the pol II mRNA, would enable Cappable-seq to also capture and identify pol II transcripts. Furthermore, by combining 5’ end Cappable-seq enrichment with 3’ polyA RNA selection would assure isolation of full-length mRNA transcripts. Coupling this with long read sequencing technologies such as SMRT sequencing (Pacific Biosciences) or Nanopore sequencing (Oxford Nanopore Technologies) would reveal the comprehensive repertoire of splice variants. In summary, by capturing the 5’ end of primary transcripts, Cappable-seq, is a profoundly unique approach to analyzing transcriptomes.

### Conclusions

Universally the initiating nucleotide found at the 5’ end of primary transcripts has a distinctive triphosphorylated end that distinguishes these transcripts from all other RNA species.

Recognizing this distinction is key to deconvoluting the primary transcriptome from the plethora of processed transcripts that confound analysis of the transcriptome. The method presented here allows for the first time capture of the 5’ end of primary transcripts. This enables a unique robust TSS determination in bacteria and microbiomes. In addition to and beyond TSS determination, Cappable-seq depletes ribosomal RNA and reduces the complexity of the transcriptome to a single quantifiable tag per transcript enabling digital profiling of gene expression in any microbiome.

## Materials and Methods

### Materials

3’ DTB-GTP synthesis was initiated with 3’-(O-Propargyl) guanosine (ChemGenes Corp. Wilmington, MA) followed by its conversion to 3’(O-Propargyl) guanosine 5’ triphosphate via a one-pot, two-step method [23]. The 3’-(O-Propargyl) Guanosine 5’ triphosphate was then purified by both ion exchange chromatography and reverse phase HPLC. The isolated 3’(O-Propargyl) guanosine 5’ triphosphate was converted to the 3’-desthiobiotin-TEG-guanosine 5’ triphosphate through the addition of desthiobiotin-TEG-azide (Berry and Associates, Inc., Dexter, MI) using copper-mediated azide-alkyne cycloaddition (“Click chemistry”, Kolb and Sharpless, Scripps Res. Inst and BaseClick, Tutzing, GmbH) [24, 25]. Final isolation of the target compound was performed using reverse phase HPLC. 2’DTB-GTP was synthesized as 3’ DTB-GTP except 2’-(O-Propargyl) guanosine was used and 3’ biotin-GTP was synthesized as 3’ DTB-GTP except that biotin-TEG-azide was substituted for desthiobiotin-TEG-azide. ATP free T4 polynucleotide kinase was prepared from T4 polynucleotide kinase (NEB) by dialysis against 10 mM Tris-HCl, 50 mM KCl, 1 mM DTT, 0.1 mM EDTA, 50% Glycerol, pH 7.4.

### Growth of *E. coli* and isolation of total RNA

*E. coli* MG1655 cells were grown at 37°C in M9 minimal media with 0.2% glucose. The culture was grown to mid-log phase and 2 volumes of RNAlater (Life Technologies) were added. The culture was incubated at 4°C overnight. The cells were collected by centrifugation and the RNA was extracted with FastRNA Blue Kit(MPBio). The RNA was then treated with DNAseI (NEB) and further purified with Megaclear kit (Life Technologies). The resulting RNA had a RIN score of 9.0 as determined by Bioanalyzer (Agilent).

### Desthiobiotin-GTP capping of *E. coli* RNA

Three micrograms of *E. coli* RNA was incubated in 50μl 1X VCE buffer (NEB) supplemented with 0.1 mM S-adenosyl methionine, and 0.5 mM DTB-GTP and 50 units of Vaccinia Capping Enzyme (NEB), for 30 minutes at 37°C. The RNA was purified on a Zymo Research Clean and Concentrator-5 column for 200 nucleotide and greater RNA per manufacturer’s instructions with a total of 4 washes with RNA wash buffer. The RNA was eluted in 100 μl of 1 mM Tris pH 7.5, 0.1 mM EDTA (low TE).

### Capture of capped T7 RNA transcript with Streptavidin

10 μl reaction volumes containing 1x VCE buffer, ^32^P uniformly labeled T7 *in vitro* 300mer transcript RNA, 10 units of VCE and either 0.5mM 2’ desthiobiotin-TEG-GTP or 3’ desthiobiotin-TEG-GTP, or GTP were incubated at 37°C for 2 hours. As carrier, 5 μl of MspI-digested pBR322 DNA (NEB) was added to the RNA and purified on MEGAclear spin columns as directed by manufacturer and eluted in 100 μl low TE. 50 μl of the eluted RNA was mixed with 50 μl of 10 mM Tris-HCl pH 7.5, 500 mM NaCl, 1mM EDTA (wash buffer A). This mix was added to the hydrophilic streptavidin magnetic beads (NEB) that had been previously prepared by washing 3 times with 400 μl of 10 mM Tris-HCl pH 7.5, 1 mM EDTA, 50 mM NaCl (wash buffer B). The beads were incubated for 10 minutes at room temperature. The beads were then washed with 100 μl of wash buffer B, and three times with 400 μl of wash buffer A, to elute unbound material. The beads were then resuspended in 50 μl of wash buffer A and an additional 50 μl of wash buffer A containing 20 mM biotin. The beads were kept resuspended for 20 minutes at room temperature by occasional quick mixing. To determine if the RNA had been selectively captured by the beads and eluted with biotin, the beads were collected on the side of the tube with a magnet, the 100 μl supernatant was collected, and radioactivity determined by scintillation counting.

### Enrichment of RNA

The desthiobiotin-GTP labeled RNA was fragmented by adding 2.5 μl of NEB 10X T4 polynucleotide kinase buffer to a 100 μl volume of capped RNA and incubated for 5 minutes at 94°C. The RNA was then collected by addition of 180 μl of AMPure XP beads plus 420 μl of 100% ethanol. The beads were washed 2X with 80% ethanol. The RNA was eluted from the beads in 100 μl of low TE. 3’ phosphates were removed from the RNA by addition 8.2 μl of 10X T4 polynucleotide buffer to 75 μl of the RNA solution and 4 μl of ATP-free T4 polynucleotide kinase (NEB) was added and incubated for 15 minutes.

Hydrophilic streptavidin magnetic beads (NEB) were prepared by washing 2 times with 400 μl of 10 mM Tris-HCl pH 7.5, 50 mM NaCl, 1 mM EDTA and 2 times with 400 μl of 10 mM Tris-HCl pH 7.5, 500 mM NaCl, 1 mM EDTA. The beads were then suspended in their original suspension concentration of 4 mg/ml in wash buffer A. 50 μl of the kinase treated RNA was added to 30 μl of the prewashed streptavidin beads at room temperature with occasional resuspension for 20 minutes. The beads were then washed two times with 200 μl of wash buffer A, and two times with 200 μl of wash buffer B. The beads were then resuspended in 30 μl of wash buffer B and 1mM biotin. The beads were incubated for 20 minutes at room temperature with occasional resuspension. The biotin eluted RNA was collected and bound to AMPure XP beads by adding 1.8 volumes of AMPure beads to the eluted RNA volume and adding 1.5 volumes of 100% ethanol to the resulting volume of the AMPure/RNA mix. The beads were washed with 80% ethanol two times and the RNA eluted with 60 μl low TE. 30 μl of the RNA eluate was added to 30 μl of prewashed streptavidin beads for a second round of enrichment. The streptavidin beads were washed and eluted as above. The biotin eluted RNA was collected and bound to AMPure beads as above and eluted with 30 μl low TE. The desthiobiotin cap was then removed to leave a 5’ monophosphate terminus by adding 3.3 μl of 10X Thermopol buffer (NEB) and 3 μl (15 units) of RppH (NEB) and incubating for 60 minutes at 37°C. The reaction was terminated by addition of 0.5 μl of 0.5 M EDTA and heating to 94°C for 2 minutes. The RNA was then bound to AMPure beads as described above, washed and eluted in 20 μl low TE.

### Mouse microbiome

Two cecum samples were obtained from two C57 female mice from which two RNA preparations were isolated. The samples were incubated in RNAlater at 4 degrees and then frozen. The RNA from the samples was prepared using Qiagen RNAeasy kit using manufacturer’s protocol. 2.4 μg of total RNA were capped with 3’DTBGTP, enriched on streptavidin beads as described above.

### RNA sequencing library prep

The NEBNext Small RNA Library Prep kit (NEB) was used to generate an Illumina sequencing library. The library was amplified through 15 cycles of PCR. RNA sequencing was performed on an Illumina MiSeq Instrument with single reads of 100 bases using V3 reagent kit. The raw reads corresponding to the *E. coli* experiment have been deposited in the European Nucleotide Archive (ENA) website under the accession number PRJEB9717, (http://www.ebi.ac.uk/ena/data/view/PRJEB9717). For the mouse microbiome analysis sequencing libraries were prepared as described above. The libraries were sequenced on an Illumina GA. Raw reads corresponding to the mouse microbiome have been deposited in the ENA website under accession number XX [submitted for accession number].

## Data analysis

### Mapping

For the *E. coli* analysis, single end reads were trimmed for adaptors using cutadapt (version 1.3) with default parameters and −a AGATCGGAAGAGCACACGTCTGAACTCCAGTCAC. The reads were mapped to the K-12 MG1655 *E. coli* genome (U00096.2) using Bowtie2 local (-L 16). To determine the 5’ end, the resulting mapped reads were trimmed to the coordinates of the most 5’ mappable end of the read (trimmed read). For the mouse microbiome analysis, all genomes in NCBI genome database from the eubacteria taxonomic group (uid 2) were downloaded. If multiple versions of the genome are available for the same species, the representative genome or reference genome was used. If no representative/reference genome were found, one version of the genome was chosen at random. Reads were trimmed for adaptors (as describe above) and mapped to each genome separately using bowtie2 with the following parameters: --local --no-1mm-upfront -L 28 --score-min G,36,17.

### Microbiome analysis

We define as present in the microbiome, bacterial species with at least 300 clustered putative TSS genome-wide. Clustered putative TSS are positions on the genome of the strongest putative TSS within 100 bp ( cluster_tss.pl --cutoff 50). A putative TSS is defined as the 5’ end position of at least one uniquely mapped read (grep -v \’XS:\’ on the mapped read sam file) using the following program: bam2firstbasegtf.pl --cutoff 0. The species with the highest number of TSS per phylum was selected as the representative species for this phylum. Next, for the representative species of each phylum, the positions of the high confident TSS were selected using the following parameters: bam2firstbasegtf.pl --cutoff 10 --absolute 1 and clustered using cluster_tss.pl --cutoff 50. This filtering resulted with 221 positions for *Lactobacillus johnsonii*, 886 positions for *Akkermansia muciniphila*, 894 positions for *Lachnospiraceae bacterium* and 174 positions for *Bifidobacterium pseudolongum* from replicate 1. For leaderless transcript annotation, the positions of the high-confident clustered TSS were compared to the annotation file for the respective species and TSS that locate at the start and in the same orientation of the annotated gene were considered as leaderless. For sequence bias analysis, the sequence context from −45 to +5 bp around the positions of the high-confident clustered TSS was compared to the overall sequence composition ([ATCG]) of the genome and a sequence logo was derived using weblogo with the following parameters: weblogo --format eps −s large −n 100 --composition [ATCG] -- yaxis 1 --errorbars NO --color-scheme classic. For read composition analysis, reads were mapped to the four representative species *(Lactobacillus johnsonii, Akkermansia muciniphila, Lachnospiraceae bacterium Bifidobacterium pseudolongum)* using Bowtie2 with the following parameters: --end-to-end --score-min ‘C,0,-1’ -L 32. The number of reads overlapping with the annotated rRNA, tRNA, coding genes and intergenic regions were computed and plotted. For the replicate analysis, high-confident clustered TSS found in either replicate 1 or replicate 2 were retained. The RRS (see below) for each retained TSS was computed in both replicate 1 and 2 for all four representative species and plotted.

### *E. coli* TSS determination

The number of trimmed reads mapping to each position on the genome is normalized to the total number of mapped reads using the following formula: *RRS = (Rns /Rt) * 1000000* with RRS being the relative read score, Rns being the number of trimmed reads mapping to position n in the *E. coli* genome on strand s (- or +) and Rt being the total number of reads mapping to the *E. coli* genome. Positions and strands with a RRS of less than 1.5 in the Cappable-seq experiment were discarded. For each of the retained positions, the RRS is compared to the RRS obtained in the control experiment using the following formula: enrichment score = log2(RRScap / *RRScontrol)* with RRScap being the RRS obtained in Cappable-seq experiment and RRScontrol being the RRS obtained in the control experiment. Positions with an enrichment score of 0 or above were considered as TSS. The suit of programs to identify, filter and cluster TSS are freely available on github (https://github.com/Ettwiller/TSS/).

### Sequence conservation for *E. coli*

Pre-computed whole genome alignments in maf format between *Escherichia coli* K12, *Escherichia coli* APEC 01, *Enterobacter* 638, *Shigella flexneri* 2a, *Salmonella typhi, Salmonella enterica* Paratypi ATCC 9150, *Yersinia pestis* CO92, *Blochmannia floridanus, Buchnera* sp. were downloaded from the UCSC microbial genome browser [26]. Conservation scores were computed using phastcon ^4^. Combining phylogenetic and hidden Markov models in biosequence analysis running phyloFit with --tree "(((((eschColi_K12,eschColi_O157H7),eschColi_APEC_O1),ente638),shigFlex_2A),(salmTyph, salmEnte_PARATYPI_ATC)yersPest_CO92)" and phastcon with the following parameters: --target-coverage 0.25 --expected-length 1. PhyloP scores were computed using the above whole genome alignment and the output of phyloFit using the following parameters: --wig-scores --method SCORE --msa-format MAF.

### *E. coli* Annotation

Gene annotations are derived from the NCBI K12 MG1665 annotation (GenBank: U00096.2). Processed sites from tRNA and rRNA are derived from the U00096.2 annotation selecting entries with feature tRNA or rRNA. The set of known TSS are derived from RegulonDB [15] (RegulonDB 8.6, 4–11–2014) combining the following files from the experimentally derived datasets: PromoterSigma24Set, PromoterSigma32Set, PromoterSigma54Set, PromoterSigma19Set, PromoterSigma28Set, PromoterSigma38Set, PromoterSigma70Set and PromoterUnknownSet. TEX comparison was done using the TSS described in supplemental file 1 (M63_0.4 condition) and table S1 *(E.coli)* from Thomason [7] and Kim [3] respectively. The composite dataset contains all the above datasets (known TSS from RegulonDB [15], Kim [3] and Thomason [7] merged into one single file).

### Comparison with TEX

Raw fastq files from the most recent d-RNA-seq experiment [7] were downloaded from ENA website accession number SRP038698. Reads were trimmed to remove the polyA tail using Trimgalor and the trimmed reads were mapped to the E. *coli* genome using bowtie local as describe above. To be in comparable conditions, the mapped reads were down-sampled to 8 millions for both TEX-, TEX +, Cappable-seq and control data.

### Motif search

Over-represented motifs were searched using MEME version 4.8.0 [27] with the -mod zoops -dna -minsites 120 -maxsize 1000000 options. Motifs logo were done using the weblogo3 program [28]).

## Data Access

The data sets supporting the results of this article are available in European Nucleotide Archive (ENA) accession number PRJEB9717, (http://www.ebi.ac.uk/ena/data/view/PRJEB9717).

## Abbreviations

bp: base pair; DTBGTP: 3’-desthiobiotin-TEG-guanosine 5’ triphosphate; R: purine; TSS: transcription start site; Y: pyrimidine; VCE: Vaccinia capping enzyme

## Authors’ contributions

IS developed the biochemistry and performed the experiments. IS and LE designed the experiments. LE performed the data analysis. JB synthesized the DTBGTP. EY assisted with library preparation and performed the sequencing. EL and IS wrote the manuscript with contributions from the other authors. All authors read and approved the final manuscript.

## Description of Additional data files

The following additional data are available with the online version of this paper. Additional file 1. Supplement. Contains all Supplementary Notes and Supplementary Figures and Supplementary Table 2. Additional file 2 contains Supplementary Table 1.

## Acknowledgments

The authors thank Nicholas Bokulich and Martin Blaser for the mouse microbiome RNA samples. We thank New England Biolabs for supporting this study.

